# Invariance of initiation mass and predictability of cell size in *Escherichia coli*

**DOI:** 10.1101/081422

**Authors:** Fangwei Si, Dongyang Li, Sarah E. Cox, John T. Sauls, Omid Azizi, Cindy Sou, Amy B. Schwartz, Michael J. Erickstad, Yonggun Jun, Xintian Li, Suckjoon Jun

## Abstract

It is generally assumed that the allocation and synthesis of total cellular resources in microorganisms are uniquely determined by the growth conditions. Adaptation to a new physiological state leads to a change in cell size via reallocation of cellular resources. However, it has not been understood how cell size is coordinated with biosynthesis and robustly adapts to physiological states. We show that cell size in *Escherichia coli* can be predicted for any steady-state condition by projecting all biosynthesis into three measurable variables representing replication initiation, replication-division cycle, and the global biosynthesis rate. These variables can be decoupled by selectively controlling their respective core biosynthesis using CRISPR interference and antibiotics, verifying our predictions that different physiological states can result in the same cell size. We performed extensive growth inhibition experiments, and discovered that cell size at replication initiation per origin, namely the initiation mass or “unit cell,” is remarkably invariant under perturbations targeting transcription, translation, ribosome content, replication kinetics, fatty acid and cell-wall synthesis, cell division, and cell shape. Based on this invariance and balanced resource allocation, we explain why the total cell size is the sum of all unit cells. These results provide an overarching framework with quantitative predictive power over cell size in bacteria.

## Introduction

Bacteria adapt their size to growth conditions. Since the 1950s, most experimental studies of the relationship between cell size and physiology have focused on nutrient conditions [1–6]. A major conclusion from these extensive studies is that cell size is coordinated with the growth rate, independent of the chemical composition of the growth medium [7]. For *E. coli,* the average cell size increases exponentially [1] and the ribosome content linearly [8,9] with respect to the nutrient-imposed growth rate. While these phenomenological “growth laws” are central to bacterial physiology, their origins have yet to be understood from more basic biological principles.

Recent studies have investigated the mechanism of cell size control via inhibition of biosynthesis [4,10]. When cells were exposed to sub-lethal dosage of a cell wall biosynthesis inhibitor, their surface-to-volume ratio changed to a new steady-state value in a dose-dependent manner [10]. These results led to a hypothesis that cell size is determined by accumulation of surface material sufficient for cell division. In an orthogonal study, inhibition of protein synthesis caused an increase or decrease in cell size depending on the inhibition method [4], implying that cell size changes are correlated with proteome reallocation by growth inhibition [9].

In principle, cell size or volume is equivalent to the total product of all biosynthesis in each division cycle. When cells are in “balanced growth,” cell size and every cellular component double at the same rate [7,11]. Therefore, a key issue for understanding cell size control is how biosynthesis of major cellular components - chromosomes, proteins, and cell envelope - is robustly coordinated with replication and division. This coordination principle should be able to predict how cell size adapts to different growth conditions. However, such a principle has not been established. Similarly, predictions of cell size for general growth conditions have not been possible, except for the special case of nutrient limitation [1].

## Results

### Deconstructing cell size control into three core biosynthetic processes

We consider a population of exponentially growing bacterial cells in steady state. The cells on average double their mass every *τ* minutes and chromosome replication initiates once per division cycle [12] (Figure 1A). One complete cell cycle, namely the combined process of chromosome replication and cell division, lasts for τ_cyc_ minutes (Figure 1A). If the cell cycle is longer than the average doubling time (τ_cyc_ > τ), the chromosome must contain multiple replication forks and duplicated replication origins (*ori*’s) (Figure 1A). The average number of *ori*’s per cell is given by 2^τ_cyc_/τ^, where τ_cyc_/τ is the average number of overlapping cell cycles (see also Supplemental Information). The power of 2 implies that the number of *ori*’s doubles when a new round of replication initiates (Figure 1A).

**Figure 1.**
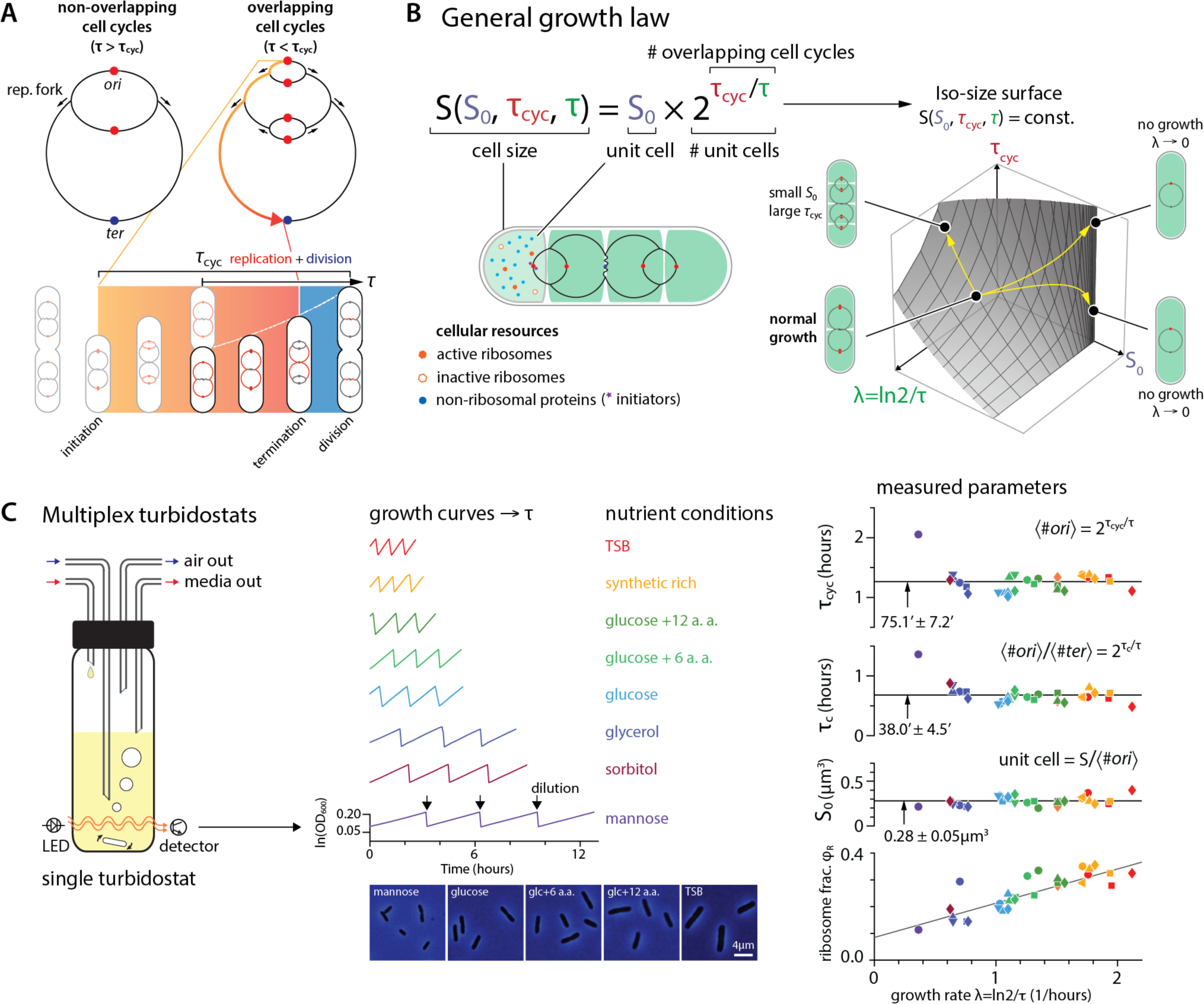
Cell cycle model and the general growth law. (A) Top: Replication of a circular chromosome from a single replication origin: non-overlapping vs. overlapping cell cycles. Bottom: Schematics to define τ_cyc_ and τ with overlapping cell cycles. (B) Left: The general growth law states that cell size is the sum of all unit cells, each unit cell containing the minimal resource for self-replication from a single replication origin. Right: If *S*_0_, τ_cyc_, and τ can vary freely and independently, there would exist an infinite number of different physiological states for the same cell size. (C) Left: Multiplex turbidostat ensures steady-state growth with automatic dilution at a pre-defined value of OD600, from which doubling time is calculated. Samples taken from each growth condition are used for imaging and cell size measurement (number of imaged cells is on the order of 10^4^; see detailed sample size in Supplemental Information). Right: Under nutrient limitation, τ_cyc_, τ_C_ and *S*_0_ remain constant. The ribosome fraction φ_R_ is measured from the same sample and shows linear increase. Symbol shapes reflect biological replicates and the colors represent different nutrient conditions. Please also see Figures S1, S6 and S7.

Formally, a simple quantitative expression relates the cell size *S* with the cell cycle duration τ_cyc_ and the doubling time *τ* as follows.

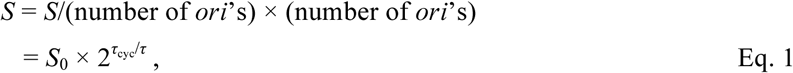

where *S*_0_ is the cell size per *ori,* proportional to the initiation mass [13]. (Here, we use size and volume interchangeably unless explicitly noted otherwise.) Conceptually, the volume *S*_0_ represents the unit cellular resources to maintain biosynthesis and start the cell cycle from one *ori* (Figure 1B). We will denote Eq. 1 the “general growth law” for its resemblance to, and generalization of (see below), the original nutrient growth law by Schaechter, Maaloe, and Kjeldaard in 1958 [1]. We will also refer the cellular resources within *S*_0_ a “unit cell,” and use the unit cell size interchangeably with the initiation mass [14]. Equation 1 suggests that the cell size is the sum of all unit cells (Figure 1B).

The general growth law formalizes the dimensional reduction of all biosynthesis into three measurable variables representing replication initiation, progression of replication-division, and the kinetics of global biosynthesis. In theory, the three variables *S*_0_, τ_cyc_, *τ* can independently vary from each other, with an infinite number of different physiological states for the same cell size (Figure 1B). The experimental and conceptual challenge is whether these three processes can actually be decoupled biologically and how they are coordinated.

### Multiplexing physiological measurements

Experimental testing of the general growth law (Eq. 1) requires extensive exploration over a large parameter space. We planned to perturb translation, transcription, DNA replication, cell division, cell wall synthesis for a wide range of growth inhibition and nutrient limitation, and quantitatively predict how cell size changes. We realized that a high-throughput single-cell approach [15,16] that led to the discovery of the “adder” principle [3,17,18] and its critical analysis [6] or the effects of growth rate fluctuations [16] was not feasible because of the large number of different experimental conditions involved. To this end, we took a population-level approach and built a multiplex turbidostat, which ensures long-term steady-state cell cultures in multiple independent growth conditions in a single experiment [19,20] (Figure 1C; see Supplemental Information). Our ultimate goal is to apply insights from population-level studies to the understanding of individual cells.

We examined the reliability of our system by reproducing known results for different nutrient conditions (Figure 1C). For each condition, we simultaneously measured cell size *S*, duration of cell cycle τ_cyc_, duration of DNA replication τ_C_, doubling time τ, and the ribosome fraction φ_R_ of the total proteome from a steady-state population (Figures 1C and S1; see Supplemental Information). The doubling time measured from OD curves is in excellent agreement with the measurements from our previous single-cell experiments in a microfluidic mother machine [15,17]. The average cell size increased exponentially with respect to the nutrient-imposed growth rate λ (= ln2/τ), in agreement with the nutrient growth law [1] (Figures 1; see Supplemental Information). The ribosome fraction φ_R_ increased linearly with the growth rate, confirming previous reports [8,9]. τ_C_ and τ_cyc_ were both constant for a wide range of growth conditions at τ_C_ = 38.0’ ± 4.5’ and τ_cyc_ = 75.1’ ± 7.2’ (Figures 1C; see Supplemental Information) [21,22]. Since τ_cyc_ is longer than the doubling time τ in these experiments, multiple cell cycles overlap. In all nutrient conditions, the unit cell size *S*_0_ remained constant, consistent with previous results that showed the constancy of the initiation mass [6,13] (Figure 1C).

### Decoupling DNA replication from growth by thymine limitation

A fundamental biological hypothesis underlying the general growth law is that the cell cycle can vary independent of growth. Testing this prediction requires an experimental means to change the duration of DNA replication without affecting the global biosynthesis rate. One possible approach is to slow replication kinetics by the reduction of nucleotides pool (Figure 2A). Following previous suggestions, we constructed a *thyA-* strain whose intracellular thymine level can be titrated by limiting thymine supplied in the growth medium [23,24]. During thymine limitation, the *thyA-* strain showed a systematic increase in τ_cyc_ by more than twofold from approximately 1 to 2.5 hours at a constant growth rate (Figure 2A).

**Figure 2.**
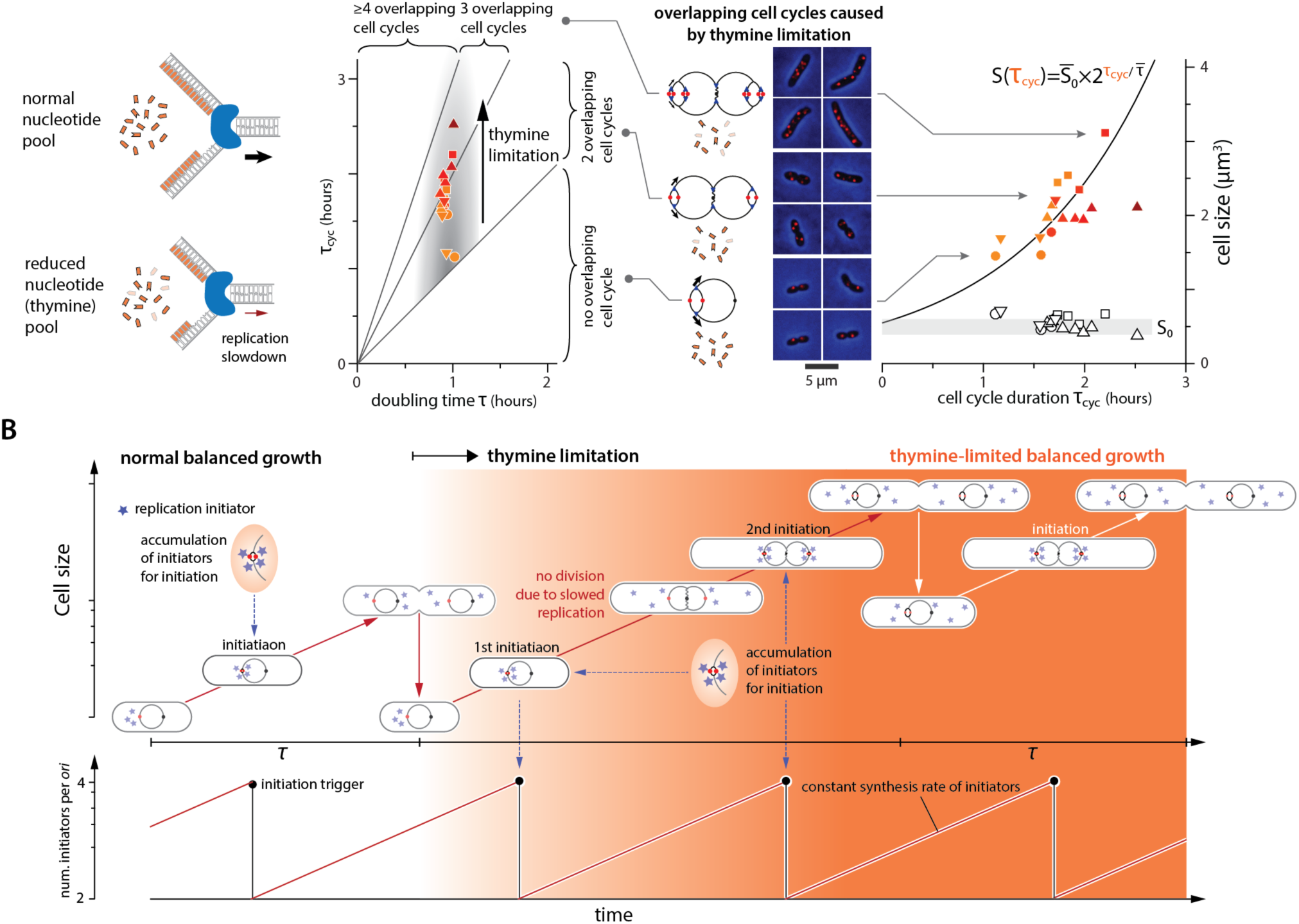
Thymine limitation alters cell cycle duration and cell size without changing the initiation mass. (A) Left: Thymine limitation reduces the nucleotide pool and replication slows consequently. Middle: τ_cyc_ increases in thymine limitation while τ remains unchanged, increasing the number of overlapping cell cycles. Chromosome schematics and cell images with foci qualitatively show increasing number of *ori*’s as a result of multifork replication. Odd number of foci in some cell images are possibly due to cohesion and/or stochasticity in replication initiation [26]. Right: Cell size increases exponentially with τ_cyc_ in thymine limitation, as predicted by Eq. 1 (solid line, no free parameters). The empty symbols are the cell size per *ori* (*S*_0_), and the thickness of the grey band denotes ±SD. Symbol shapes reflect biological replicates and the symbol colors indicate the level of thymine limitation. (B) Explanation of increasing cell size in thymine limitation. Thymine limitation is applied at the beginning of the second generation, and replication slows. Initiation-competent initiators accumulate at the same rate as the growth rate λ and trigger initiation at a critical number per *ori* (four in this illustration). An extra round of replication is initiated during transition as cell division is delayed due to slowed replication. Cell size reaches new steady state in the third generation. Bottom panel shows constant rate of accumulation of initiation-competent replication initiators. Please also see Figure S2.

The prolonged replication period had non-trivial consequences on the cell cycle. Since the doubling time remained constant at τ *≈* 1 hour, the 2.5-fold increase in τ_cyc_ caused up to three overlapping cell cycles. Direct visualization of the *ori* region using ParS-ParB-mCherry [25,26] further confirmed multifork replication in *thyA-*cells (Figure 2A).

These observations contrast with the classic *E. coli* cell cycle model based on nutrient limitation, in which only fast growing cells exhibit multifork replication [27]. In thymine limitation, even slow-growing cells can overlap multiple cell cycles due to increased τ_cyc_ (Figure 2A).

Coordination between size, growth, and the cell cycle under thymine limitation can be explained based on a class of models that a critical number of initiation-competent initiators per *ori* must accumulate to trigger initiation (which we generically refer to as the “initiator threshold” model) (Figure 2B) [28–30]. Upon thymine limitation, replication rate decreases and replication cannot finish by the time the cell would have divided under the normal conditions. In the meantime, initiators continue to accumulate at the same rate as the constant growth rate, and can reinitiate another round of replication while division is being delayed. After completing the initial round of replication, the cell divides at a larger size due to the delayed division, and the newborn cells contain chromosomes that are partially replicated. Consequently, cell size increases exponentially with increased τ_cyc_ in thymine limitation, whereas the growth rate, ribosome fraction and the unit cell size remain constant, quantitatively supporting the general growth law (Figures 2A and S2A).

### Decoupling τ_cyc_ using CRISPR interference

Our thymine limitation results show that it is possible to change the cell size in a quantitatively predictive manner by selective control of the core biosynthesis represented in the general growth law. To test the decoupling hypothesis further using a genetic method, we developed tunable CRISPR interference (“tCRISPRi”), a plasmid-free CRISPRdCas9 based suppression system [20]. tCRISPRi allows precise and continuous titration of gene expression by more than 10-fold from the wild-type expression level (Figures 3A and 3B).

**Figure 3.**
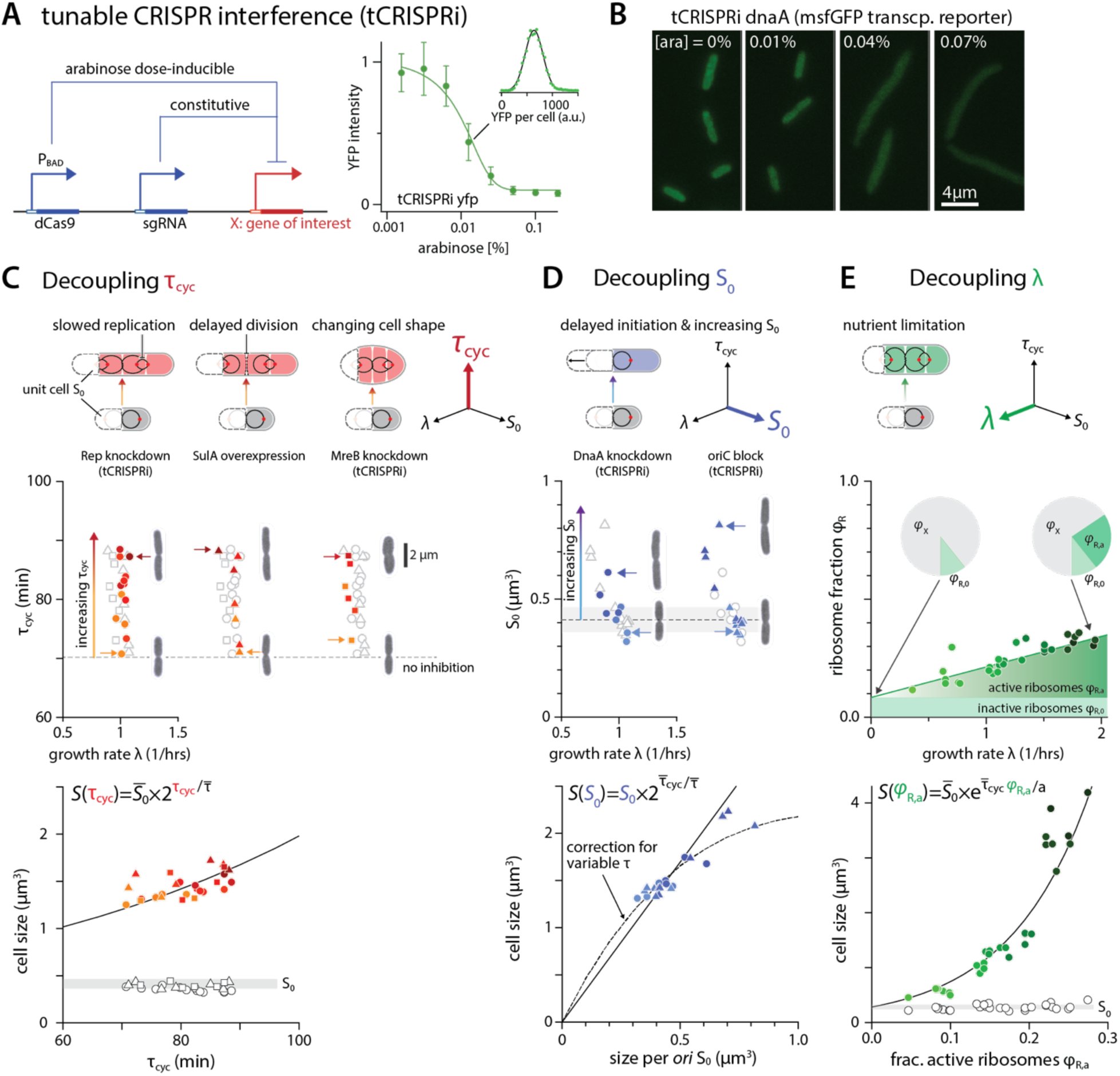
Decoupling of τ_cyc_, *S*_0_ and λ by selective inhibition of biosynthesis using tunable CRISPR interference. (A) In the tCRISPRi strain, dCas9 is induced from an engineered P_BAD_ promoter in a dose-dependent manner by arabinose, repressing the targeted gene with the help of specific sgRNA. Constitutive YFP is knocked down to demonstrate the tunability of the system. Figures are adapted from [20]. (B) Example images from tCRISPRi DnaA-expressing strain. The co-transcriptional reporter msfGFP level decreases as DnaA is knocked down. (C) Top: τ_cyc_ can be decoupled from *S*_0_ and λ by three orthogonal methods: slowing replication (Rep knockdown; circles, left), cell division (SulA over-expression; triangles, middle), or changing cell shape (MreB knockdown; squares, right). The symbol colors represent the degree of knockdown or overexpression (same for Figures 3D and 3E). Bottom: Cell size increases exponentially as predicted by the general growth law (solid line, no adjustable parameters; same for Figures 3D and 3E). The unit cell size *S*_0_ remains unchanged (open symbols). Grey band indicates average *S*_0_ from no-induction controls and its thickness indicates ±SD. (D) Top: The unit cell size *S*_0_ can be decoupled from τ_cyc_ and λ using two orthogonal methods: repression of DnaA (filled circles, left) or sequestration of *oriC* (filled triangles, right). Bottom: Cell size increases following the general growth law. The solid line is Eq. 1 with constant λ. The dashed line is Eq. 1, assuming a linear dependence of λ on *S*_0_ (fitted separately) to account for the slight decrease in growth rate in the *S*_0_ vs. λ data. Grey band indicates average *S*_0_ from no-induction controls and its thickness indicates ±SD. (E) Top: Decoupling λ from τ_cyc_ and *S*_0_ by nutrient limitation. Bottom: The nutrient growth law, namely the exponential dependence of the average size on λ, is a special case of Eq. 1, where *S*_0_ and τ_cyc_ are constant. The growth rate can be expressed in terms of the active ribosome fraction φ_R,a_. The unit cell size *S*_0_ is constant over all growth rates. Grey band indicates the average *S*_0_ with thickness indicating ±SD. Please also see Figure S3.

Using tCRISPRi, we first set out to decouple DNA replication from growth. Inactivating proteins in the replisome was an obvious alternative to thymine limitation, and we made a tCRISPRi strain to repress DNA helicase Rep because its mutation is known to slow replication forks [31]. As expected, the replication period increased about twofold during Rep knockdown, while the growth rate remained constant (Figures 3C and S3D). In addition, we found that Rep knockdown does not cause morphological changes, a major side effect of thymine limitation (Figure 2A). As an independent test, we exposed cells to hydroxyurea, which mildly inhibits ribonucleotide reductase and slows DNA replication as well [32]. The results were consistent with Rep knockdown (Figure S3I).

These observations contrast with the classic *E. coli* cell cycle model based on nutrient limitation, in which only fast growing cells exhibit multifork replication [27]. In thymine limitation, even slow-growing cells can overlap multiple cell cycles due to increased τ_cyc_ (Figure 2A).

Coordination between size, growth, and the cell cycle under thymine limitation can be explained based on a class of models that a critical number of initiation-competent initiators per *ori* must accumulate to trigger initiation (which we generically refer to as the “initiator threshold” model) (Figure 2B) [28–30]. Upon thymine limitation, replication rate decreases and replication cannot finish by the time the cell would have divided under the normal conditions. In the meantime, initiators continue to accumulate at the same rate as the constant growth rate, and can reinitiate another round of replication while division is being delayed. After completing the initial round of replication, the cell divides at a larger size due to the delayed division, and the newborn cells contain chromosomes that are partially replicated. Consequently, cell size increases exponentially with increased τ_cyc_ in thymine limitation, whereas the growth rate, ribosome fraction and the unit cell size remain constant, quantitatively supporting the general growth law (Figures 2A and S2A).

### Decoupling τ_cyc_ using CRISPR interference

Our thymine limitation results show that it is possible to change the cell size in a quantitatively predictive manner by selective control of the core biosynthesis represented in the general growth law. To test the decoupling hypothesis further using a genetic method, we developed tunable CRISPR interference (“tCRISPRi”), a plasmid-free CRISPRdCas9 based suppression system [20]. tCRISPRi allows precise and continuous titration of gene expression by more than 10-fold from the wild-type expression level (Figures 3A and 3B).

Using tCRISPRi, we first set out to decouple DNA replication from growth. Inactivating proteins in the replisome was an obvious alternative to thymine limitation, and we made a tCRISPRi strain to repress DNA helicase Rep because its mutation is known to slow replication forks [31]. As expected, the replication period increased about twofold during Rep knockdown, while the growth rate remained constant (Figures 3C and S3D). In addition, we found that Rep knockdown does not cause morphological changes, a major side effect of thymine limitation (Figure 2A). As an independent test, we exposed cells to hydroxyurea, which mildly inhibits ribonucleotide reductase and slows DNA replication as well [32]. The results were consistent with Rep knockdown (Figure S3I).

A notable feature in the general growth law is the symmetry between the replication period and the division period in τ_cyc_. In other words, Eq. 1 predicts that slowing replication and delaying cell division have identical effects on cell size via τ_cyc_. We tested this prediction by delaying cell division with SulA overexpression [33] or cephalexin treatment (Figures 3C, S3F and S3H). Both methods increased τ_cyc_ selectively, similar to the increase in τ_cyc_ by replication slowdown.

Additionally, we found that morphological change by MreB depletion also decouples τ_cyc_ via a delay in cell division [34,35]. MreB is a major cytoskeletal protein and we found its knockdown by tCRISPRi causes a gradual morphological transition from rod to round shape. τ_cyc_ increased monotonically, consistent with Rep knockdown or SulA overexpression. Yet, both the unit cell and the growth rate remained unchanged (Figures 3C and S3E).

In these decoupling experiments, the cell size increased exponentially with respect to τ_cyc_ following the general growth law (Figures 3C, S3D, S3E, S3F, S3H and S3I). Taken together with thymine limitation, our results reaffirm it is possible to decouple chromosome replication and cell division from growth, and manipulate cell size in a controlled manner.

### Decoupling the initiation mass by initiation delay

Next, we sought to decouple the initiation mass from growth and τ_cyc_. Conceivably, initiation mass would change if the initiation timing is selectively perturbed without affecting global biosynthesis. We tested these ideas using two tCRISPRi strains to repress a major replication initiator protein DnaA [36,37] and sequester the initiation site *oriC* [38], respectively. In both experiments, the initiation mass *S_0_* increased systematically in a dose-dependent manner, and cell size increased linearly with the unit cell size *S*_0_ as predicted by Eq. 1 (Figures 3D, S3A and S3C).

### Decoupling growth and the origin of the nutrient growth law

The general growth law encompasses the nutrient growth law [1] and explains its origin. Under nutrient limitation, average cell size is exclusively determined by the growth rate. However, our decoupling experiments show that the duration of cell cycle τ_cyc_ and the initiation mass *S*_0_ can also vary by selective inhibition of biosynthesis (Figures 3C and 3D). The nutrient growth law is therefore a special case of the general growth law, wherein the growth rate is the only experimental variable, decoupled from **τ**_cyc_ and *S*_0_ by nutrient limitation [1].

The mechanism of growth rate control is currently unknown. Nevertheless, since the fraction of active ribosomes φ_R,a_ of the total proteome is a proxy of global biosynthesis, growth rate may be directly proportional to φ_R,a_ under nutrient limitation (Figure 3E) [9,39]. This leads to an exponential dependence of cell size on φ_R,a_ by the general growth law (Figure 3E).

### Invariance of initiation mass under extensive perturbations to global biosynthesis

Our results show that growth and the cell cycle can be decoupled by targeted inhibition of biosynthesis, and cell size changes following the general growth law. A natural question is to what extent the general growth law can predict cell size when cells are exposed to global physiological perturbations.

We examined the predictive power of Eq. 1 by applying a broad pallet of antibiotics targeting different biosynthesis. We observed significant deviation in cell size vs. growth rate from the nutrient growth law, without any obvious pattern (Figure 4A). In stark contrast to the changes in cell size and growth rate, the unit cell size *S_0_* remained invariant under extensive perturbations targeting transcription, translation, ribosome content, fatty acid synthesis, cell wall synthesis, replication speed or cell division (Figure 4B). The distribution of *S*_0_ was a tight Gaussian with an average of 0.28 ± 0.05 μm^3^. This matches the extrapolated cell size (0.27 ± 0.07 μm^3^) in the nutrient growth law as the growth rate approaches zero (Figure 4B).

**Figure 4.**
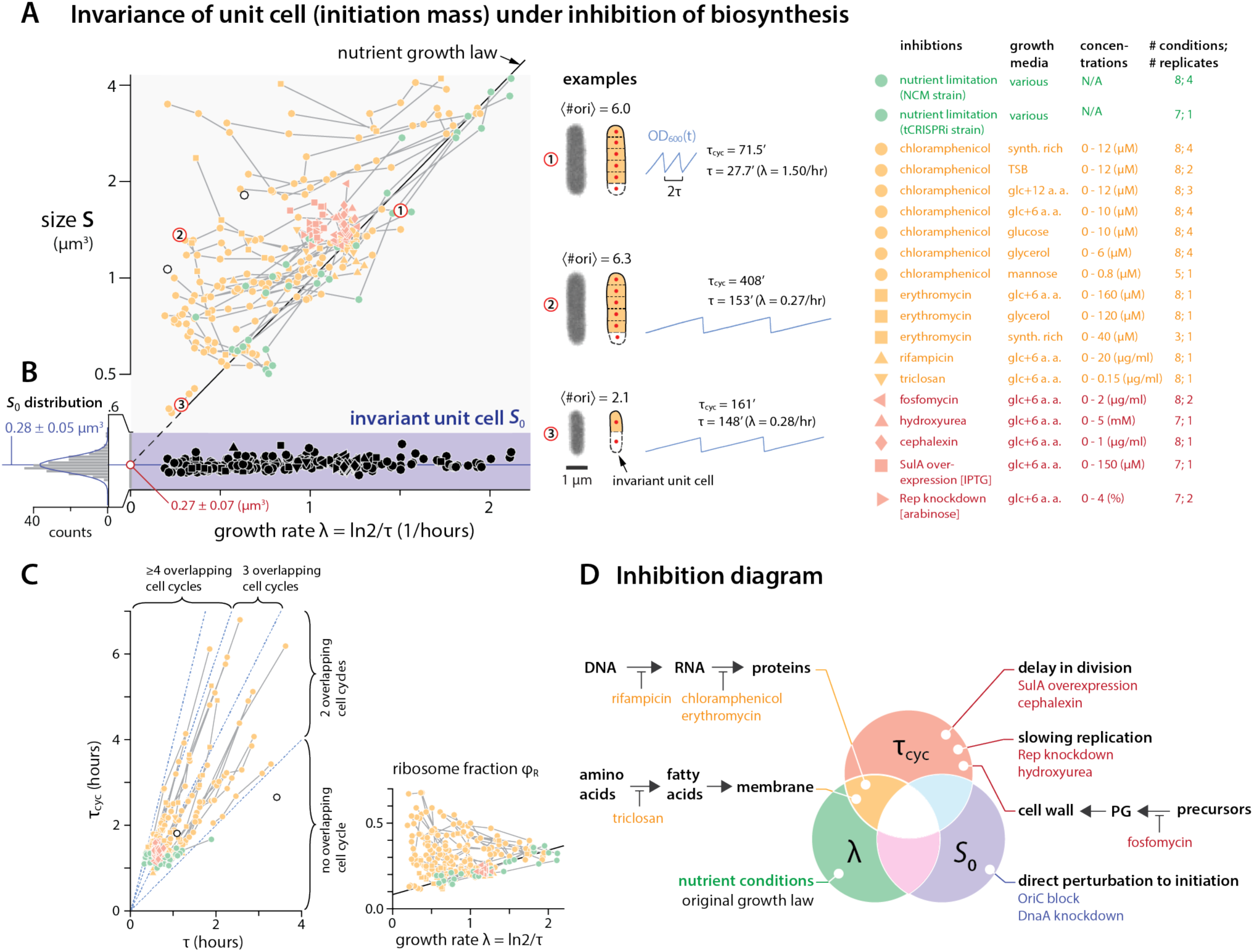
Invariance of unit cell under extensive growth inhibition. (A) Cell size versus growth rate extensively deviates from the nutrient growth law (see legend to the right for experimental conditions). Numbered circles show three exemplary data points. The two empty circles represent pooled single-cell data from [6]. Coloring reflects which core biosynthetic process is perturbed (same for Figures 4C and 4D). (B) The initiation mass or unit cell size *S_0_* remains invariant despite extensive changes in average size and growth rate. The distribution of *S*_0_ is well fitted by Gaussian, and its average (0.28 ± 0.05 μm^3^) coincides with the y-intercept of the nutrient growth law (0.27 ± 0.07 μm^3^). (C) Left: The measured τ_cyc_ vs. τ shows a linear relationship under growth inhibition. Right: The ribosome fraction φ_R_ increases when the growth rate decreases by growth inhibition. Empty circles represent pooled single-cell data from [6]. (D) An "inhibition diagram" mapping perturbations to the three core biosynthetic processes underlying the general growth law. Please also see Figures S2 and S4.

The invariance of the unit cell was unexpected, because transcriptional and translational inhibition significantly affected the cell cycle (Figure 4C, left; Figures S2 and S4). In particular, the average doubling time τ and the cell cycle duration τ_cyc_ increased linearly proportional to each other, similar to nutrient limitation in slow growth conditions [4,6,21] (Figures 4C, left). Furthermore, the proteome partitioning measured by an RNA/protein ratio was also altered significantly by inhibition of global biosynthesis (Figure 4C, right) [9]. Therefore, these results suggest that the invariant unit cell under the inhibition of global biosynthesis is a basic, invariant building block of cell size.

Based on the decoupling results (Figure 3), we classify physiological perturbations in terms of how they reallocate cellular resource and change the three core biosynthetic processes underlying the general growth law (Figure 4D; see Discussion).

### Universal structure of cell size control and coordination of biosynthesis

A universal property of cell size control emerges from our data. Under extensive perturbations, cell size vs. growth rate in general do not follow the exponential relationship of the nutrient growth law (Figure 4A, left and Figure 5A). However, when the cell size and the doubling time are rescaled by their respective *S*_0_ and τ_cyc_, the rescaled size *S/S*_0_ vs. growth rate τ_cyc_/τ collapses onto a single exponential master curve (Figure 5B). This again underscores that the exponential relationship between size and growth rate in the original nutrient growth law [1] is a special case of the general growth law where *τ* is the only experimental variable (Figures 3E and 5A). Equation 1 is also a good approximation for single cells in steady-state growth, and recent single-cell data [6] collapse reasonably well onto the same master curve (Figures 5B and S5).

**Figure 5.**
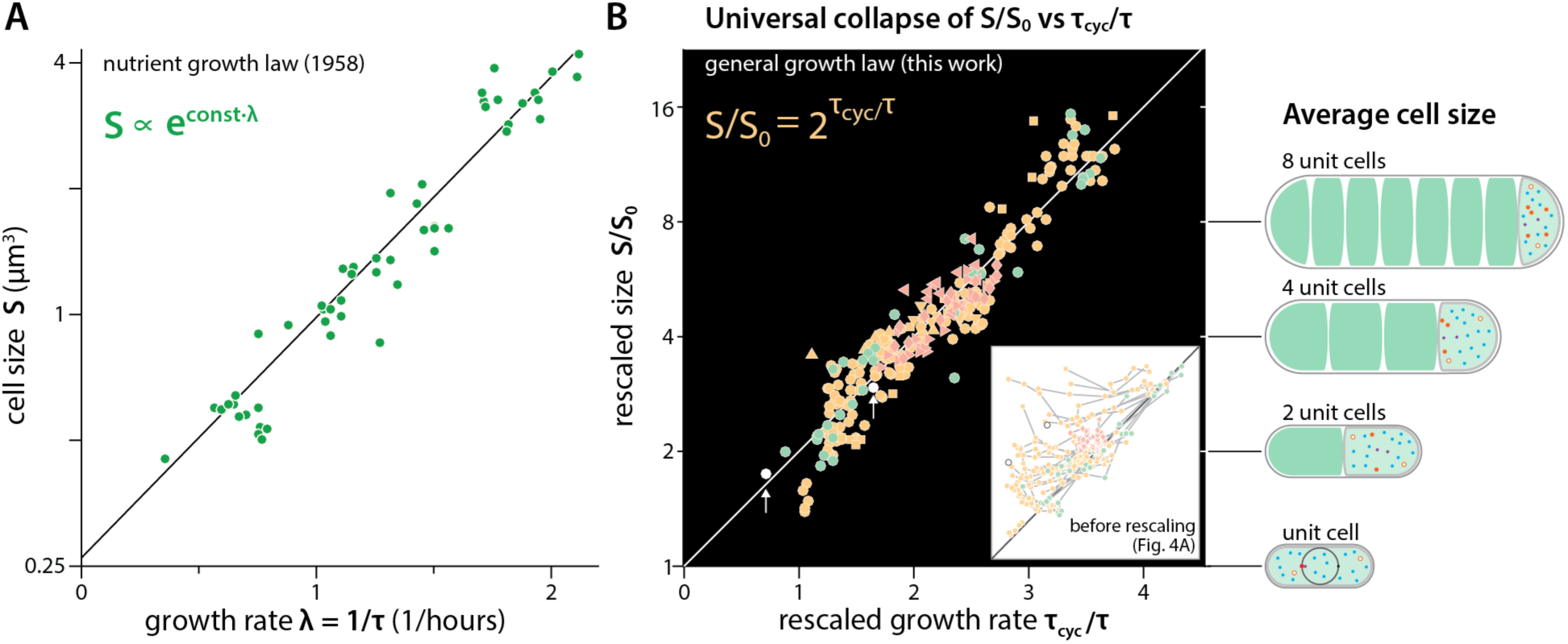
From the nutrient growth law (1958) to the general growth law. (A) The nutrient growth law by Schaechter, Maaløe, and Kjeldgaard (1958) [1] prescribes an exponential relationship between average cell size *S* and growth rate λ under nutrient limitation. Data points are taken from this study. (B) The general growth law extends the nutrient growth law. Not only λ but also *S*_0_ and τ_cyc_ are experimental variables. All raw data from Figure 4 (inset) collapse onto a single master curve after rescaling, demonstrating the predictive power of the general growth law and the origin of the nutrient growth law. Empty circles (with arrows) represent the average of pooled single-cell data from [6]. The cell diagrams on the right illustrate that the average cell size is the sum of all unit cells. Please also see Figure S5.

The universal collapse demonstrates that all biosynthesis can be mapped onto three coarse-grained variables *S*_0_, τ_cyc_, τ (Figure 4D), and their coordination abides by the general growth law (Eq. 1), allowing prediction of cell size for any steady-state condition.

## Discussion

The general growth law provides a simple and straightforward prescription for cell size. It states that cell size is the sum of all unit cells for any steady-state growth condition. Despite its apparent simplicity, the general growth law and its meaning was previously underappreciated. The reason is partly historical in that most previous studies on cell size and physiology focused on understanding the nutrient growth law [1,40] and cell cycle control under different nutrient conditions.

The invariance of the unit cell under growth inhibition significantly extends the concept of constant initiation mass which was limited to varying nutrient conditions [6,13]. While the mechanism of the invariance is unknown, recent single-cell data have provided an important insight into this constancy [6]. In particular, the initiation mass for individual cells was constant in two different nutrient conditions, independent of the birth size and the elongation rate in individual cells. This is consistent with the initiator threshold model [27,28,30] that we used to explain the thymine limitation results (Figure 2). In support of the model, the concentration of a key initiator protein DnaA is known to be constant under different nutrient conditions [41]. Perhaps more important, the ratio of its ATP and ADP forms remained constant even when DnaA is significantly overexpressed [42].

To explain the invariance of the unit cell, the initiator threshold model alone is not sufficient; the initiator concentration must also be invariant independent of the growth condition and growth inhibition [43,44]. Therefore, our results predict the existence of a specific protein “sector” that remains constant for a wide range of physiological perturbations that changes the ribosome fraction of the proteome [4, 9] (Figure 6B). If the replication initiators indeed belong to this hypothetical sector, invariance of the unit cell size is ensured by balanced growth as illustrated in Figure 6A despite growth inhibition.

**Figure 6.**
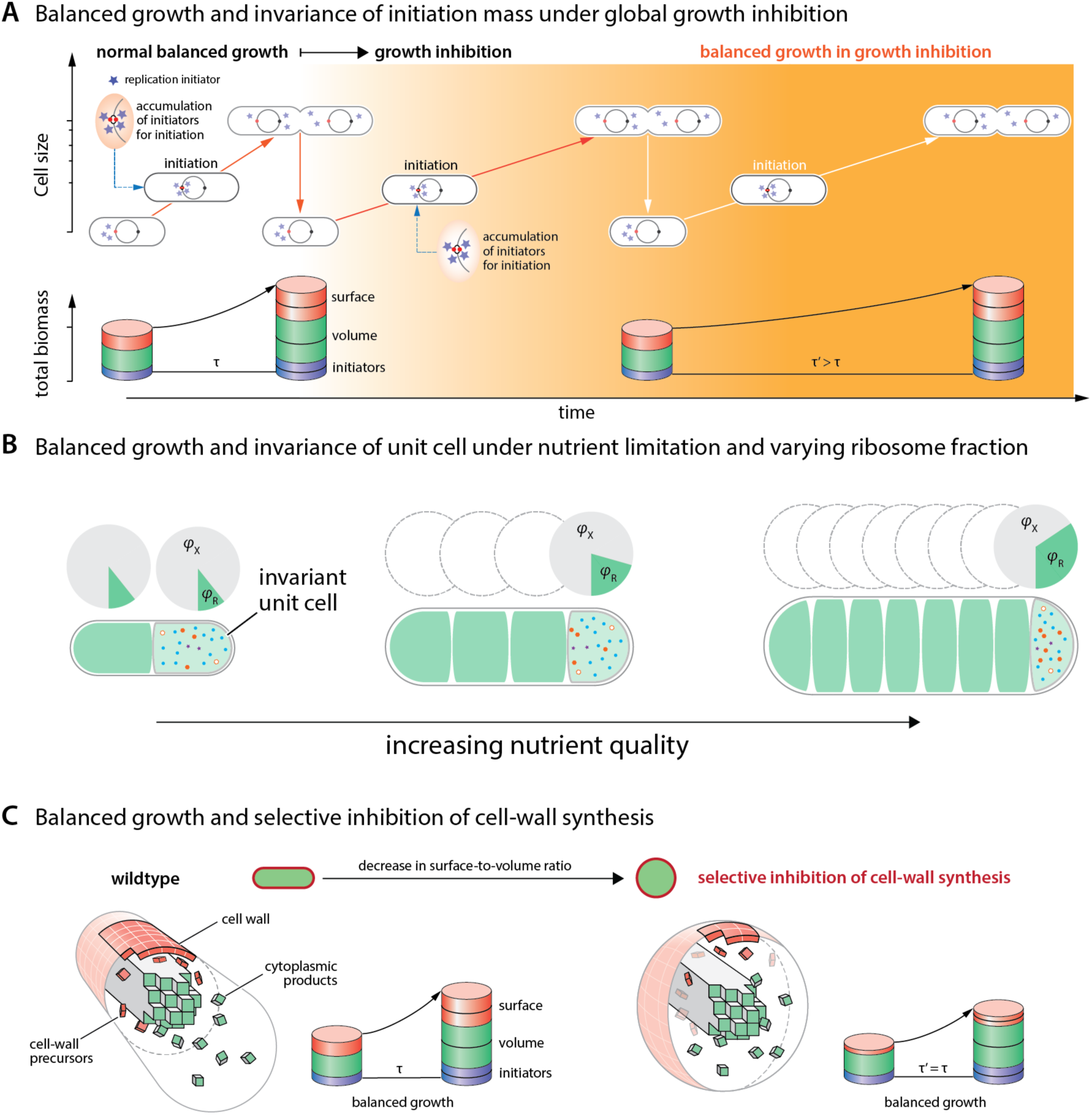
Invariance of initiation mass and resource allocation explain cell size in balanced growth. (A) Inhibition of global biosynthesis decreases the synthesis rate of all intracellular components by the same degree, without changing their concentrations or the cell cycle dynamics. Therefore, the initiation mass remains invariant. (B) In nutrient limitation, ribosome fraction changes linearly with respect to growth rate. However, the initiation mass remains invariant due to a constant fraction of “initiator” sector, independent of changes in the rest of proteome fraction. Under nutrient limitation, ribosome fraction increases with the growth rate, causing an increase in cell size. (C) Inhibition of cell-wall synthesis rebalances the ratio of available materials for surface growth and volume growth. Cells must reduce their surface-to-volume ratio to accommodate more volumetric biomass per unit surface area, and thus become more round. The cell size changes due to a new steady-state value of τ_cyc_ in the general growth law, without changing *S_0_* or λ.

When combined with the invariance of the unit cell, the general growth law helps understand which cell cycle stage is responsible for changes in cell size under selective inhibition of biosynthesis. For example, recent work and our data show that treatment of fosfomycin causes *E. coli* and other bacterial organisms to reduce their surface-to-volume ratio and increase their size (Figures 6C and S3G) [10]. When cell-wall synthesis is selectively inhibited, relatively less surface materials are produced compared to the cytoplasmic products, and cells must become more round to adapt to the reduced amounts of cell-wall materials (Figure 6C). Since the initiation mass, duration of replication, and growth rate remained unchanged in our experiments, the sole source of the size change is the cell division kinetics of the round cell (Figure S3G; see Supplemental Information). Similar quantitative analysis allows for prediction of τ_cyc_ changes in LacZ overexpression experiments (Figure S4M) [4].

Considering the minimal assumptions underlying Eq. 1, the general growth law has obvious implications on other microorganisms.

## Experimental Procedures

### Strains and growth conditions

All experiments in normal growth conditions or with physiological perturbations were carried out using NCM 3722 [45] unless otherwise noted. Thymine limitation and tCRISPRi experiments were based on strains with a MG1655 background [20]. Detailed information of strain genotypes is included in Supplemental Information. Before every turbidostat experiment, cells were inoculated into 1 ml lysogeny broth (LB) medium as seed culture from a single colony on agar plate, streaked no more than 7 days before use. After 6-12 hours in 30°C or 37°C water bath shaker, cells were diluted 1000-fold into 1 ml - 2 ml of the specific growth medium as pre-culture and shaken at 37°C in water bath until OD_600_ = 0.2. The pre-culture was then back-diluted 1000-fold again into the same medium and shaken at 37°C in water bath until OD_600_ = 0.2. The back-diluted culture was then inoculated into each turbidostat vial with or without specific inhibiting conditions, and the turbidostat experiment was started with OD_600_ ≈ 0.05 for each vial. The turbidostat was then run for at least 8 generations in steady-state growth before sample collection. In each turbidostat experiment, we were able to run up to 8 different growth conditions simultaneously. For each condition, samples were collected for cell size, RNA/protein and cell cycle measurements. See Supplemental Information for experimental details.

### Growth rate measurement

To ensure exponential growth, cells were kept between OD_600_ = 0.05 and OD_600_ = 0.20. Once the OD_600_ reaches 0.20, the vial was diluted automatically until OD_600_ = 0.05 (Figure 1C). The OD vs. time curve was fit to a single exponential *I* = *I_0_* 2^t/τ^ for each segment between two consecutive dilution events and the average was taken as the generation time. Growth rate is given by λ=ln2/τ. Each turbidostat is blanked with the same media before each experiment. For calibration of OD from the phototransistors, see Supplemental Information for details.

### Microscopy and image acquisition

All transmission light and fluorescence microscopy were performed on an inverted microscope (Nikon Ti-E) with Perfect Focus 2 (PFS 2), 100x oil immersion objective (PH3, NA=1.45), LED fluorescent light (Lumencor Inc., OR), and Andor NEO sCMOS camera (Andor Technology Ltd., MA). Exposure time was between 100-200ms with 100% transmission. From each set of sample, 140-300 images were captured and 5,000-30,000 cells were analyzed to ensure statistically significant distributions of cell measurements such as cell size or DNA content. All cell image analysis was carried out by custom software written in Python employing the OpenCV library.

### Cell size measurement

Cell samples were fixed with 0.24% w/v formaldehyde in the same growth media and imaged within 24 hours. Image analysis extracted the contours of all possible items from phase contrast images, and these were primarily filtered based on shape and size. The probability distribution of contour width is symmetrical and not correlated with cell length [17]. All filtered contours within 3 standard deviations from the mean value of the distributions were kept and exported as isolated images for re-examination. Cell size as well as surface area were calculated as a cylinder with two hemispherical ends for rod-shaped cells. The aspect ratio was calculated as the ratio of length to width of cell contour.

### RNA/protein measurements

Total protein was measured as previously reported [46] except for the following modifications for sample collection. For some nutrient conditions, cells were collected when OD_600_=0.2, so 6 ml cell culture was collected. Total RNA was measured as previously reported [46] except for the following modifications for sample collection. For some nutrient conditions, cells were collected when OD_600_=0.2, so 6 ml cell culture was collected. The cells were pelleted by centrifugation, washed once with 1 ml water, centrifuged again and the pellet fast frozen on dry ice.

### Cell cycle measurement (qPCR & Image cytometry)

C period was estimated by marker frequency analysis using qPCR. Genomic DNA was prepared from the turbidostat sample and amplified using PowerUp SYBR Green Master Mix (ABI). Quantification of DNA was done by ∆Ct method as previously reported [47]. A total of 8 pairs of primers targeting different chromosomal loci were used. The ratio of relative copy numbers of two loci gives the ratio of C period over generation time (C/τ) as 〈ori〉/〈ter〉=2^C/τ^ (see Supplemental Information). C period was then calculated using the generation time as discussed above. DNA content was measured by image cytometry instead of flow cytometry [48]. Ethanol-fixed samples were stained with 3μg/ml Hoechst 33342 [32]. Standard cells with known DNA content were mixed with sample cells to calibrate the absolute DNA content. See details in Supplemental Information.

### Statistics and data analysis

No statistical methods were used to predetermine sample size, and the dataset table of full measurement is provided together with the Supplemental Information. The sample size N for each experiment is typically on the order of 10^4^. Exact numbers can be found in supplemental tables (‘Growth conditions and sample size’). The standard errors [sdev/sqrt(N)] are smaller than the data symbols due to the large sample sizes. For each experimental condition, we obtained 2-4 biological replicates. The systematic errors in doubling time and cell size measurements due to experimental setup or data analysis methods are smaller than the stochasticity of those variables measured from single-cell study [17].

## Author Contributions

F.S, D.L., S.C., O.A., C.S., A.S., Y.J., X.L. conducted experiments. J.T.S. and M.E. designed and built the turbidostats. F.S, D.L., S.C., J.T.S., O.A, S.J. analyzed data. S.J. designed the research. F.S., D.L., and S.J. wrote the paper.

## Acknowledgements

This work was supported by the Paul G. Allen Family Foundation, Pew Charitable Trust, NSF CAREER, and NIH R01 GM118565-01 (to S.J.). We are deeply grateful to Arshad Desai, Johan Paulsson, James Pelletier, Matt Scott, Terry Hwa, Tsutomu Katayama, Petra Levin, Kirsten Skarstad, Massimo Vergassola, and all members of Jun lab for critical reading and invaluable discussions. Special thanks to Tony Hui (Princeton) for help with RNA/protein measurements in the initial stage of this work, and Don Court for helpful experimental suggestions.

## References

1. Schaechter, M., Maaløe, O., and Kjeldgaard, N.O. (1958). Dependency on medium and temperature of cell size and chemical composition during balanced growth of Salmonella typhimurium. Microbiology 19, 592–606.

2. Hill, N.S., Kadoya, R., Chattoraj, D.K., and Levin, P.A. (2012). Cell size and the initiation of DNA replication in bacteria. PLoS Genet. 8, e1002549.

3. Campos, M., Surovtsev, I. V., Kato, S., Paintdakhi, A., Beltran, B., Ebmeier, S.E., and Jacobs-Wagner, C. (2014). A Constant Size Extension Drives Bacterial Cell Size Homeostasis. Cell 159, 1433–1446.

4. Basan, M., Zhu, M., Dai, X., Warren, M., Sévin, D., Wang, Y.P., and Hwa, T. (2015). Inflating bacterial cells by increased protein synthesis. Mol. Syst. Biol. 11, 836.

5. Kennard, A.S., Osella, M., Javer, A., Grilli, J., Nghe, P., Tans, S.J., Cicuta, P., and Cosentino Lagomarsino, M. (2016). Individuality and universality in the growth-division laws of single *E. coli* cells. Phys. Rev. E 93, 12408.

6. Wallden, M., Fange, D., Lundius, E.G., Baltekin, Ö., and Elf, J. (2016). The Synchronization of Replication and Division Cycles in Individual *E. coli* Cells. Cell 166, 729–739.

7. Neidhardt, F.C. (1999). Bacterial growth: constant obsession with dN/dt. J. Bacteriol. 181, 7405–7408.

8. Bremer, H., and Dennis, P.P. (2008). Modulation of Chemical Composition and Other Parameters of the Cell at Different Exponential Growth Rates. EcoSal Plus 3.

9. Scott, M., Gunderson, C.W., Mateescu, E.M., Zhang, Z., and Hwa, T. (2010). Interdependence of cell growth and gene expression: origins and consequences. Science 330, 1099–1102.

10. Harris, L.K., and Theriot, J.A. (2016). Relative Rates of Surface and Volume Synthesis Set Bacterial Cell Size. Cell 165, 1479–1492.

11. Campbell, A. (1957). Synchronization of cell division. Bacteriol. Rev. 21, 263–272.

12. Mitchison, J.M. (1971). The biology of the cell cycle (Cambridge University Press).

13. Donachie, W.D. (1968). Relationship between cell size and time of initiation of DNA replication. Nature 219, 1077–1079.

14. Donachie, W.D., and Begg, K.J. (1970). Growth of the bacterial cell. Nature 227, 1220–1224.

15. Wang, P., Robert, L., Pelletier, J., Dang, W.L., Taddei, F., Wright, A., and Jun, S. (2010). Robust growth of *Escherichia coli*. Curr. Biol. 20, 1099–1103.

16. Hashimoto, M., Nozoe, T., Nakaoka, H., Okura, R., Akiyoshi, S., Kaneko, K., Kussell, E., and Wakamoto, Y. (2016). Noise-driven growth rate gain in clonal cellular populations. Proc. Natl. Acad. Sci. U. S. A. 113, 3251–3256.

17. Taheri-Araghi, S., Bradde, S., Sauls, J.T., Hill, N.S., Levin, P.A., Paulsson, J., Vergassola, M., and Jun, S. (2015). Cell-Size Control and Homeostasis in Bacteria. Curr. Biol. 25, 1–7.

18. Jun, S., and Taheri-Araghi, S. (2015). Cell-size maintenance: universal strategy revealed. Trends Microbiol. 23, 4–6.

19. Toprak, E., Veres, A., Yildiz, S., Pedraza, J.M., Chait, R., Paulsson, J., and Kishony, R. (2013). Building a morbidostat: an automated continuous-culture device for studying bacterial drug resistance under dynamically sustained drug inhibition. Nat. Protoc. 8, 555–567.

20. Li, X., Jun, Y., Erickstad, M.J., Brown, S.D., Parks, A., Court, D.L., and Jun, S. (2016). tCRISPRi: tunable and reversible, one-step control of gene expression. Sci. Rep. 6, 39076.

21. Helmstetter, C. (1996). Timing of Synthetic Activities in the Cell Cycle. In Escherichia coli and Salmonella: Cellular and Molecular Biology, F. C. Neidhardt and R. Curtiss, eds. (ASM Press), pp. 1627–1639.

22. Stokke, C., Flåtten, I., and Skarstad, K. (2012). An Easy-To-Use Simulation Program Demonstrates Variations in Bacterial Cell Cycle Parameters Depending on Medium and Temperature. PLoS One 7, e30981.

23. Pritchard, R.H., and Zaritsky, A. (1970). Effect of thymine concentration on the replication velocity of DNA in a thymineless mutant of *Escherichia coli* Nature 226, 126–131.

24. Zaritsky, A., and Pritchard, R.H. (1973). Changes in Cell Size and Shape Associated with Changes in the Replication Time of the Chromosome of *Escherichia coli*. J. Bacteriol. 114, 824–837.

25. Nielsen, H.J., Ottesen, J.R., Youngren, B., Austin, S.J., and Hansen, F.G. (2006). The *Escherichia coli* chromosome is organized with the left and right chromosome arms in separate cell halves. Mol. Microbiol. 62, 331–338.

26. Youngren, B., Nielsen, H.J., Jun, S., and Austin, S. (2014) The multifork *Escherichia coli* chromosome is a self-duplicating and self-segregating thermodynamic ring polymer. Gene Dev. 28, 71–84.

27. Helmstetter, C.E., and Cooper, S. (1968). DNA synthesis during the division cycle of rapidly growing *Escherichia coli* B/r. J. Mol. Biol. 31, 507–518.

28. Fantes, P.A., Grant, W.D., Pritchard, R.H., Sudbery, P.E., and Wheals, A.E. (1975). The regulation of cell size and the control of mitosis. J. Theor. Biol. 50, 213–244.

29. Helmstetter, C., Cooper, S., Pierucci, O., and Revelas, E. (1968). On the bacterial life sequence. Cold Spring Harb. Symp. Quant. Biol. 33, 809–22.

30. Donachie, W.D., and Blakely, G.W. (2003). Coupling the initiation of chromosome replication to cell size in *Escherichia coli*. Curr. Opin. Microbiol. 6, 146–150.

31. Lane, H.E., and Denhardt, D.T. (1975). The rep mutation. IV. Slower movement of replication forks in *Escherichia coli* rep strains. J. Mol. Biol. 97, 99–112.

32. Odsbu, I., Morigen, and Skarstad, K. (2009). A reduction in ribonucleotide reductase activity slows down the chromosome replication fork but does not change its localization. PLoS One 4, e7617.

33. de Boer, P.A., Crossley, R.E., and Rothfield, L.I. (1990). Central role for the *Escherichia coli* minC gene product in two different cell division-inhibition systems. Proc. Natl. Acad. Sci. 87, 1129–1133.

34. Peters, J.M., Colavin, A., Shi, H., Czarny, T.L., Larson, M.H., Wong, S., Hawkins, J.S., Lu, C.H.S., Koo, B.-M., Marta, E., et al. (2016). A Comprehensive, CRISPR-based Functional Analysis of Essential Genes in Bacteria. Cell 165, 1493–1506.

35. Zheng, H., Ho, P.-Y., Jiang, M., Tang, B., Liu, W., Li, D., Yu, X., Kleckner, N.E., Amir, A., and Liu, C. (2016). Interrogating the *Escherichia coli* cell cycle by cell dimension perturbations. Proc. Natl. Acad. Sci. 113, 15000–15005.

36. Løbner-Olesen, A., Skarstad, K., and Hansen, F.G. (1989). The DnaA protein determines the initiation mass of *Escherichia coli* K-12. Cell 57, 881–889.

37. Skarstad, K., and Katayama, T. (2013). Regulating DNA replication in bacteria. Cold Spring Harb Perspect Biol 5, 1–17.

38. Wiktor, J., Lesterlin, C., Sherratt, D.J., and Dekker, C. (2016). CRISPR-mediated control of the bacterial initiation of replication. Nucleic Acids Res. 44, 3801–3810.

39. Scott, M., Klumpp, S., Mateescu, E.M., and Hwa, T. (2014). Emergence of robust growth laws from optimal regulation of ribosome synthesis. Mol. Syst. Biol. 10, 747.

40. Klumpp, S., Zhang, Z., and Hwa, T. (2009). Growth Rate-Dependent Global Effects on Gene Expression in Bacteria. Cell 139, 1366–1375.

41. Hansen, F.G., Atlung, T., Braun, R.E., Wright, A., Hughes, P., and Kohiyama, M. (1991). Initiator (DnaA) protein concentration as a function of growth rate in *Escherichia coli* and Salmonella typhimurium. J. Bacteriol. 173, 5194–5199.

42. Kurokawa, K., Nishida, S., Emoto, a, Sekimizu, K., and Katayama, T. (1999). Replication cycle-coordinated change of the adenine nucleotide-bound forms of DnaA protein in *Escherichia coli*. EMBO J. 18, 6642–6652.

43. Sompayrac, L., and Maaløe, O. (1973). Autorepressor Model for Control of DNA Replication. Nature 241, 133–135.

44. Hansen, F.G., Christensen, B.B., and Atlung, T. (1991). The initiator titration model: computer simulation of chromosome and minichromosome control. Res. Microbiol. 142, 161–167.

45. Brown, S.D., and Jun, S. (2015). Complete Genome Sequence of *Escherichia coli* NCM3722. Genome Announc. 3, e00879–15.

46. Hui, S., Silverman, J.M., Chen, S.S., Erickson, D.W., Basan, M., Wang, J., Hwa, T., and Williamson, J.R. (2015). Quantitative proteomic analysis reveals a simple strategy of global resource allocation in bacteria. Mol. Syst. Biol. 11, 784–e784.

47. Schmittgen, T.D., and Livak, K.J. (2008). Analyzing real-time PCR data by the comparative CT method. Nat. Protoc. 3, 1101–1108.

48. Vischer, N.O.E., Huls, P.G., Ghauharali, R.I., Brakenhoff, G.J., Nanninga, N., and Woldringh, C.L. (1999). Image cytometric method for quantifying the relative amount of DNA in bacterial nucleoids using *Escherichia coli*. J. Microsc. 196, 61–68.

